# Precursor processing by SBT3.8 and phytosulfokine signaling contribute to drought stress tolerance in Arabidopsis

**DOI:** 10.1101/2020.10.21.349779

**Authors:** Nils Stührwohldt, Eric Bühler, Margret Sauter, Andreas Schaller

## Abstract

Increasing drought stress poses a severe threat to agricultural productivity. Plants, however, evolved numerous mechanisms to cope with such environmental stress. Here we report that the stress-induced production of a peptide signal contributes to stress tolerance. The expression of phytosulfokine (PSK) peptide precursor genes, and transcripts of three subtilisin-like serine proteases, SBT1.4, SBT3.7 and SBT3.8 were found to be up-regulated in response to osmotic stress. Stress symptoms were enhanced in *sbt3.8* loss-of-function mutants and could be alleviated by PSK treatment. Osmotic stress tolerance was improved in plants overexpression the precursor of PSK1 (*proPSK1*) or *SBT3.8* resulting in higher fresh weight and improved lateral root development in the transgenic compared to wild-type plants. We further showed that SBT3.8 is involved in the biogenesis of the bioactive PSK peptide. ProPSK1 was cleaved by SBT3.8 at the C-terminus of the PSK pentapeptide. Processing by SBT3.8 depended on the aspartic acid residue adjacent to the cleavage site. ProPSK1 processing was impaired in the *sbt3.8* mutant. The data suggest that increased expression in response to osmotic stress followed by the post-translational processing of proPSK1 by SBT3.8 leads to the production of PSK as a peptide signal for stress mitigation.

**Highlight:** The expression of phytosulfokine precursor genes and processing by the subtilase SBT3.8 are upregulated in response to osmotic stress for improved drought tolerance in Arabidopsis.

## Introduction

Plants have evolved mechanisms to adapt to their environment and are thus able to survive unfavorable stress conditions. Due to climate change, the ability to cope with increasing temperatures and water scarcity becomes more and more important. Plant responses to these abiotic stress factors are regulated at multiple levels including peptide signals (Takahashi and Shinozaki, 2019)

Several classes of plant signaling peptides are distinguished based on differences in structure and biogenesis. Some peptides are derived from larger precursor proteins that may or may not have a function on their own. Other peptides are produced from small peptide-encoding genes, or translated from short open-reading frames within the 5’-untranslated regions of protein-coding genes, or from the primary transcript of miRNAs (Tavormina et al., 2015). Precursor-derived peptides can be divided into cysteine-rich peptides that depend on disulfide bond formation for activity, and the large group of post-translationally modified peptides (Matsubayashi, 2014). The latter require proteolytic processing to release the peptide moiety from the inactive precursor, and may rely on tyrosine sulfation, proline hydroxylation and arabinosylation of hydroxyprolines for full activity (Stührwohldt and Schaller, 2019).

Recent publications reported that peptide precursor processing is essential for the biogenesis of mature and active peptides, and that the processing activity may be the limiting factor for peptide signaling. IDA (Inflorescence Deficient in Abscission) is processed by subtilisin-like serine proteases (SBTs) SBT4.12, SBT4.13 and SBT5.2 and plants lacking SBT activity are impaired in floral organ abscission (Schardon et al., 2016). The tyrosine-sulfated CLEL6 and CLEL9 peptides are activated by several steps of proteolytic processing that occur in consecutive compartments of the secretory pathway (Stührwohldt et al., 2020). Following cleavage of the signal peptide upon entry into the Endoplasmic Reticulum (ER), CLEL6 and 9 are pre-processed by SBT6.1 in the *cis*-Golgi and activated by aspartate-dependent SBTs including SBT3.8 in post Golgi compartments (Stührwohldt et al., 2020; Ghorbani et al., 2016). Processing and release of the PEP1 peptide as an immune-modulatory phytocytokine are regulated by calcium-dependent activation of metacaspase 4 (Hander et al., 2019). Formation of the embryonic cuticle is controlled by the SBT2.4 (ALE1)-dependent activation of Twisted Seed 1 (TWS1; (Doll et al., 2020)), and the drought-induced formation of Phytosulfokine (PSK) by phytaspase 2 regulates flower drop in tomato (Reichardt et al., 2020)

In addition to drought-induced tomato flower drop, the disulfated PSK pentapeptide (Y(SO_3_H)-I-Y(SO_3_H)-T-Q) is involved also in the regulation of several developmental processes and biotic as well as abiotic interactions. Briefly, PSK promotes growth of roots and hypocotyls mainly by regulation of cell expansion by a plasma-membrane localized module including leucine-rich repeat receptor kinases PSKR1 and PSKR2, the co-receptor BRI1-associated receptor kinase 1 BAK1, the proton pump AHA1 and the cyclic nucleotide-gated channel CNGC17 (Stührwohldt et al., 2011; Kutschmar et al., 2009; Matsubayashi et al., 2006; Ladwig et al., 2015). Furthermore in *Zinnia elegans*, PSK promotes the differentiation of mesophyll cells into tracheary elements by down-regulation of stress-response genes like chitinases, redox-related enzymes or ethylene synthesis genes at the onset of the trans-differentiation process (Matsubayashi et al., 1999; Motose et al., 2009). Defense responses against biotrophic and necrotrophic are affected differentially by PSK signalling. Arabidopsis plants deficient in PSK receptor genes are more susceptible to pathogens with a necrotrophic lifestyle whereas resistance against hemi- or biotrophs is increased (Loivamäki et al., 2010; Igarashi et al., 2012; Mosher et al., 2013; Rodiuc et al., 2016). Also in tomato, loss of PSK receptor function led to increased susceptibility to the necrotroph *Botrytis cinerea* (Zhang et al., 2018). In contrast, PSK attenuates pattern-triggered immunity in Arabidopsis resulting in increased resistance of the PSK receptor mutant against *Pseudomonas syringae* (Igarashi et al., 2012)

PSK depends on tyrosine sulfation for bioactivity. Tyrosine sulfation is driven by Golgi-localized tyrosylprotein sulfotransferase (TPST) that is encoded by a single-copy gene in Arabidopsis (Komori et al., 2009). The enzymes that are responsible for precursor processing to release the PSK peptide are essentially unknown in Arabidopsis. SBT1.1 was shown to cleave Arabidopsis PSK4 precursor peptides *in vitro* (Srivastava et al., 2008), but the cleavage site is upstream of the mature peptide within the variable part of the precursor, and the physiological relevance of this cleavage and of SBT1.1 for PSK biogenesis remain unclear. In tomato, an aspartate-specific SBT (tomato phytaspase 2, SlPhyt2) was shown to be involved in PSK maturation (Reichardt et al., 2020). Indeed all eight PSK precursors in tomato (Reichardt et al., 2020), and the seven PSK precursors in Arabidopsis (Kaufmann and Sauter, 2019) share an aspartate residue on the amino side of the PSK pentapeptide. Arabidopsis proPSK1 is the only precursor featuring aspartate at both, the N- and the C-terminal processing sites.

Inspired by the finding that PSK mediates drought-induced flower drop in tomato (Reichardt et al., 2020) we investigated whether PSK and SBT-mediated PSK maturation are also involved in drought stress responses in Arabidopsis. Because drought stress is difficult to standardize under experimental conditions, mannitol-induced water scarcity is commonly used as an experimental approximation to drought (Hazman et al., 2016). In this paper we identified PSK genes and SBTs that are up-regulated upon mannitol-treatment. We identify SBT3.8 from Arabidopsis as the limiting factor in the osmotic stress response and the formation of PSK from the PSK propeptide. We describe a novel mechanism of peptide-dependent adaptation to osmotic stress that relies on SBT3.8-mediated release of PSK as the stress signal.

## Results

To address a potential involvement of SBTs and PSK signaling in drought stress responses in Arabidopsis, the regulation of *SBT* and *PSK* gene expression was analyzed in response to osmotic stress. Seedlings were grown for five days on control media and then transferred to mannitol-containing media. Gene expression was compared in seedlings exposed to osmotic stress and control seedlings by qPCR. We observed that three out of 56 *SBT* genes (*SBT1.4, SBT3.7*, and *SBT3.8*), and three out of five *PSK* genes (*PSK1, PSK3*, and *PSK4*) were up-regulated in response to mannitol-treatment (Figure 1). Levels of *SBT* induction were similar to genes previously shown to be regulated under osmotic stress conditions (Sham et al., 2015) including *At3g14067* (which is the *SBT1.4* gene), *At3g46280* and *At2g42540* (Figure 1).

**Figure 1:**
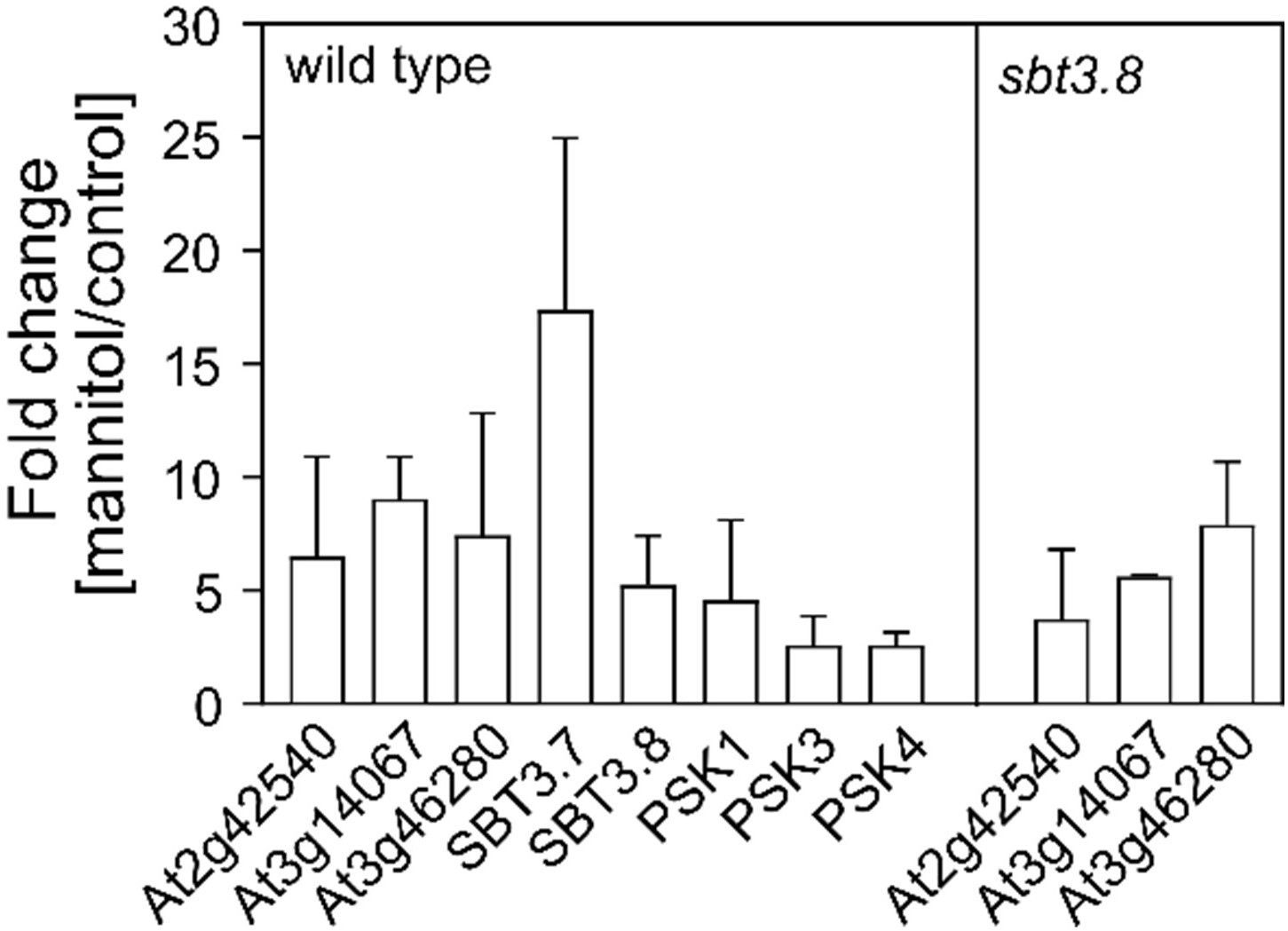
*SBT1.4, SBT3.7, SBT3.8* and PSK precursor genes are up-regulated upon osmotic stress treatment. Relative expression of *At2g42540, At3g14067* (alias *SBT1.4*), *At3g46280, SBT3.7, SBT3.8, PSK1, PSK3* and *PSK4* in wild type and in the *sb3.8* loss-of-function mutant plants upon mannitol treatment. Plants were grown for 5 days under control conditions and transferred to plates containing no or 300 mM of mannitol for six additional days. qPCR was performed on three biological replicates with two technical repeats, and gene regulation was normalized to three reference genes. Each time point included pooled plant material of several independent plants.

Next we analyzed osmotic stress tolerance of *sbt1.4, sbt3.7*, and *sbt3.8 loss-of-function* mutants. While the *sbt1.4* and *sbt3.7* mutants did not seem to be affected (data not shown) *sbt3.8* was clearly more sensitive to mannitol treatment than wild type (Figure 2). The reduction in shoot and root fresh weight was significantly higher for the *sbt3.8* mutant compared to wild type (Figure 2B, C). At the age of 5 days, when seedlings were transferred from control media to mannitol-containing media they did not have any lateral roots. The number of lateral roots produced during six days on mannitol-containing medium was significantly lower for the *sbt3.8* loss-of-function mutant compared to wild type (Figure 2D). Impaired lateral root development, and more severe reduction of shoot and root growth of the *sbt3.8* mutant suggest a role for SBT3.8 in the acclimation to osmotic stress. Despite the apparent reduction in stress tolerance, osmotic stress markers were induced in the *sbt3.8* mutant to levels comparable to wild type (Figure 1). Therefore, SBT3.8 does not seem to be required for osmotic stress perception, but may rather be involved in downstream signaling events.

**Figure 2:**
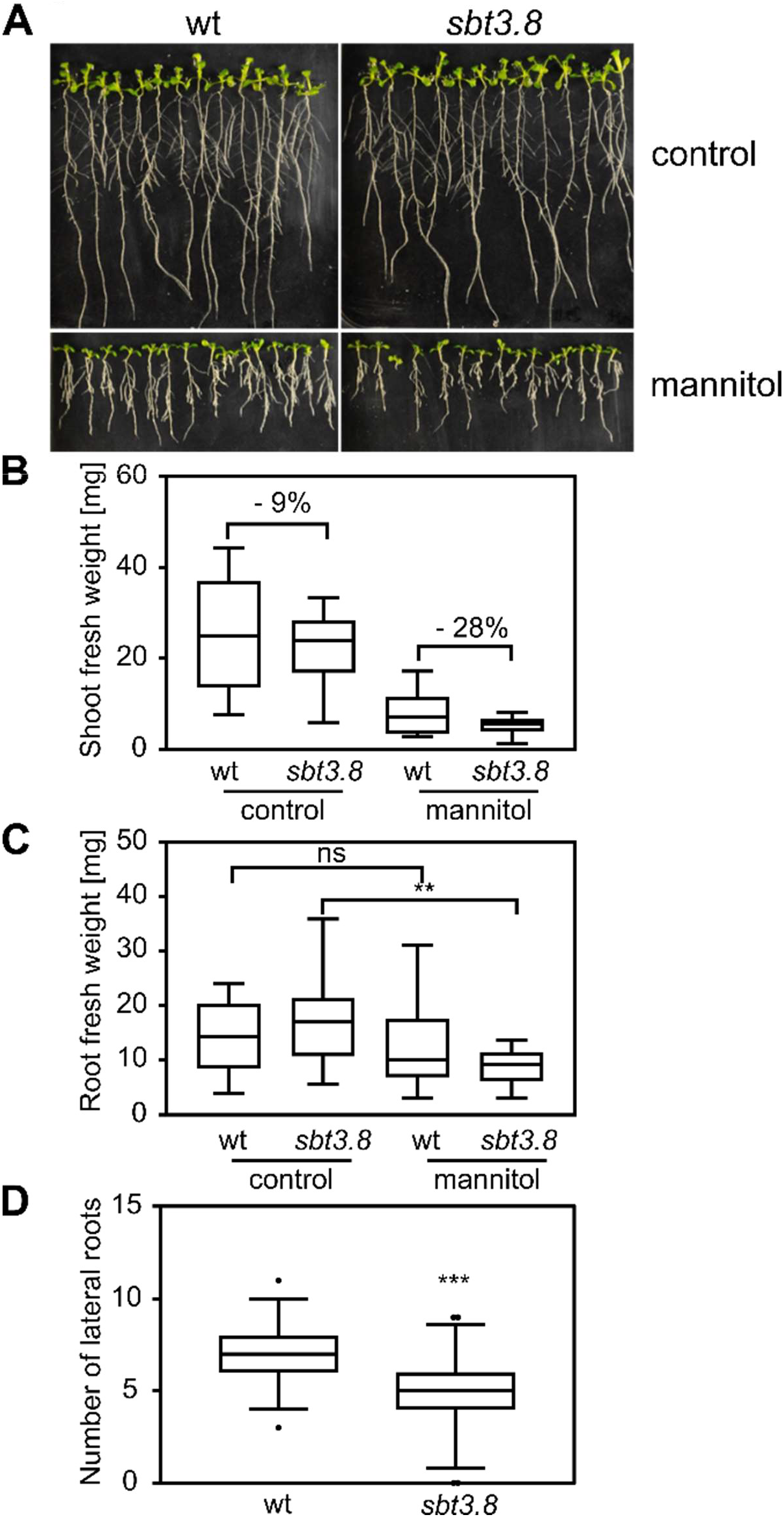
*sbt3.8* loss-of-function plants are more sensitive to osmotic stress. (A) Representative wild-type (wt) and *sbt3.8* plants grown under control conditions or with 300 mM mannitol. (B-C) Shoot (B) and root (C) fresh weight [mg] of wt and *sbt3.8* plants grown for 5 days under control conditions and transferred to plates containing no or 300 mM of mannitol for six additional days. (D) Number of lateral roots of wt and *sbt3.8*. Asterisks show significant differences to the indicated condition or genotype. Data are shown for one representative of at least three independent experiments as the mean ± SE ((B) n ≥ 13, (C) n ≥ 11, (D) n ≥ 47). *** and ** indicate significant differences at P < 0.001 or P < 0.01, respectively (two-tailed t test).

As PSK is known to promote root growth and lateral root development (Kutschmar et al., 2009), and since three out of seven PSK genes were up-regulated upon drought/osmotic stress treatment, we applied the PSK peptide in the seedling growth bioassay to assess a potential contribution of PSK to stress tolerance. Again, *sbt3.8* mutants were more sensitive to mannitol treatment, showing a stronger reduction in root and shoot growth than wild type (Figure 3A, middle). Growth of both wt and *sbt3.8* seedlings was improved upon addition of PSK (Figure 3A, bottom) consistent with a role for PSK in stress mitigation. Again, we quantified the number of lateral roots after transfer to media containing mannitol. The number of lateral roots was induced by PSK in both wild type and *sbt3.8*. However, the PSK-induced increase in lateral root number was much higher in *sbt3.8* (+ 59 %) compared to wild type (+ 15 %, Figure 3B), suggesting that endogenous levels of PSK may be reduced in *sbt3.8*. The data are thus consistent with impaired PSK formation in the *sbt3.8* mutant.

**Figure 3:**
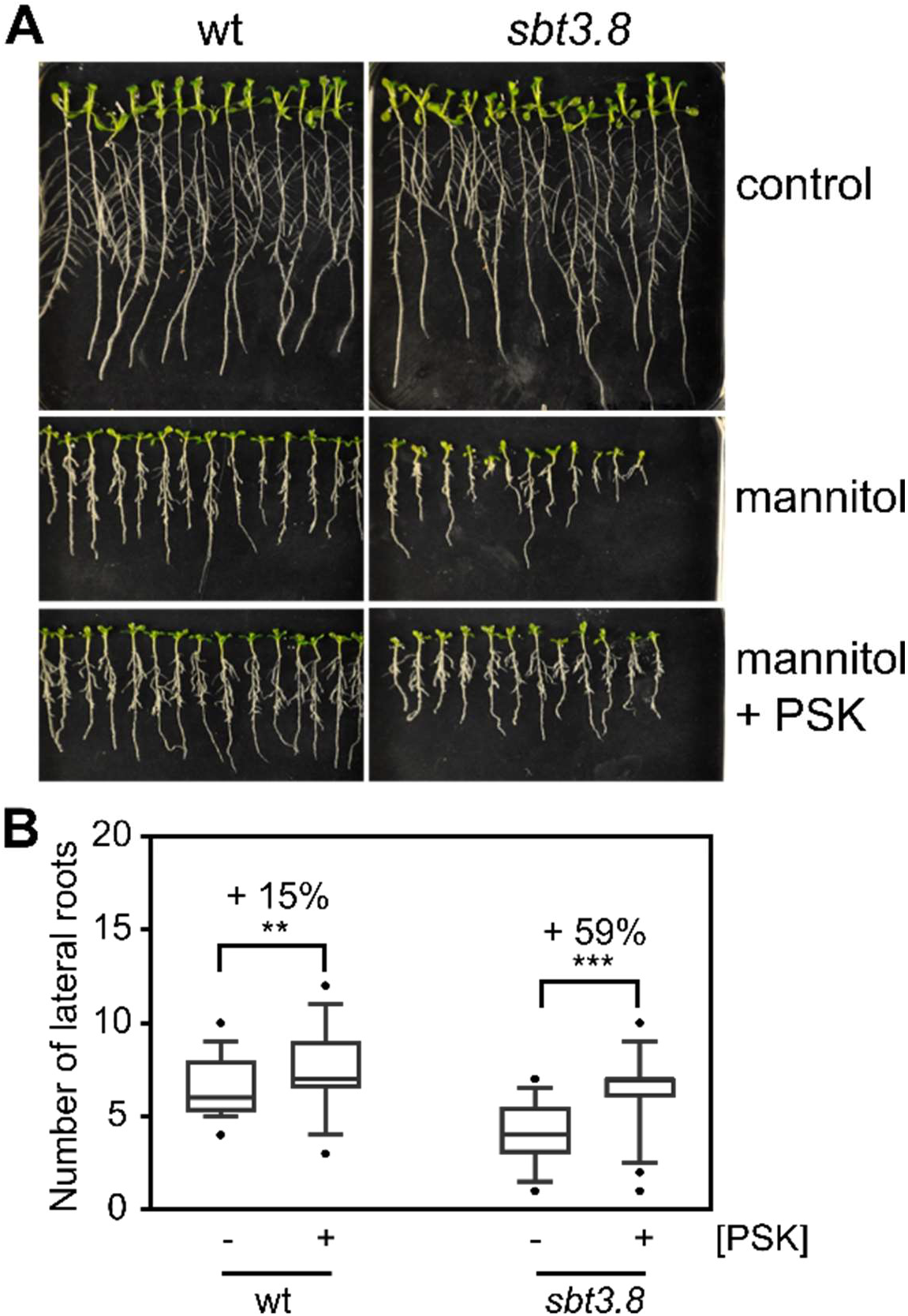
PSK can recover the mannitol-induced *sbt3.8* loss-of-function phenotype. (A) Representative pictures of wild-type (wt) and *sbt3.8* plants grown under control conditions, with 300 mM mannitol, or with 300 mM mannitol and 1 μM PSK. (B) Number of lateral roots of wt and *sbt3.8* plants grown as in (A). Data are shown for one representative experiment as the mean ± SE (n ≥ 49)). *** and ** indicate significant differences at P < 0.001 or P < 0.01, respectively (two-tailed t test).

PSK formation requires at least two steps of proteolytic processing to release the bioactive pentapeptide from its precursor. The *PSK* gene most strongly up-regulated in response to osmotic stress was *PSK1* (Figure 1). In contrast to other PSK precursors, in which the PSK pentapeptide is flanked by aspartic acid (Asp, D) only at the N-terminus, proPSK1 features Asp at both, the N- and the C-terminal processing sites. Asp is located immediately upstream of the scissile bond (in P1) at the N-terminal site, and downstream of the scissile bond (in P1’) at the C-terminus (Figure 4A). Interestingly, SBT3.8 has been described as an Asp-specific protease (a phytaspase; (Chichkova et al., 2018)). SBT3.8 was found to be highly selective for Asp at acidic pH (pH 5.5), cleaving a panel of synthetic peptide substrates with Asp in P1 (Chichkova et al., 2018), and the CLEL6 precursor with Asp in P1’ (Stührwohldt et al., 2020). We thus tested whether proPSK1 is a substrate of SBT3.8, and if so, which site is cleaved by SBT3.8.

**Figure 4:**
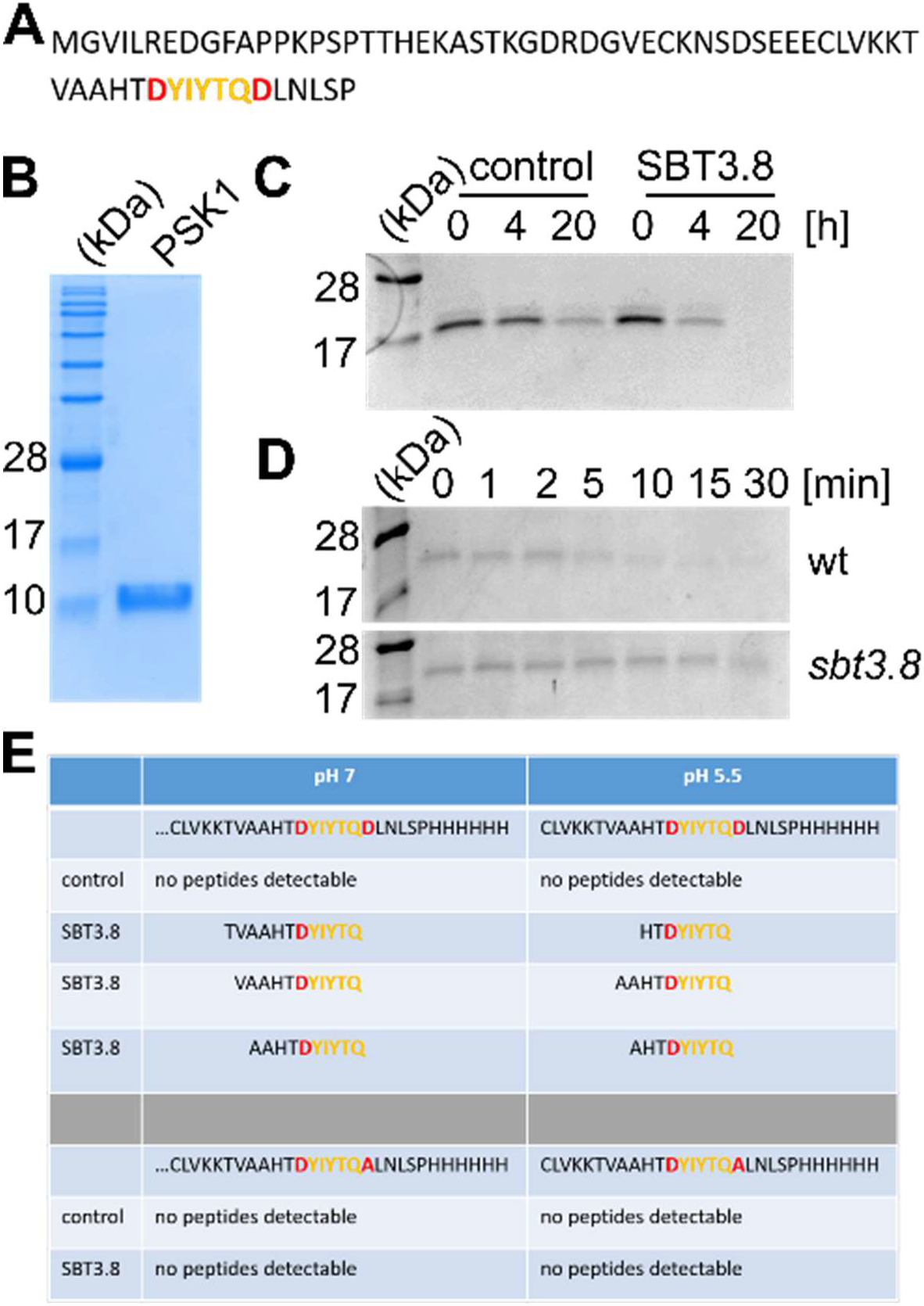
C-terminal processing of the PSK precursor by SBT3.8 depends on Asp in P1’. (A) Sequence of the wild type PSK1 precursor expressed in *E. coli*. Red letters show aspartates flanking the PSK pentapeptide (YIYTQ, orange) in P1 at the N-terminal, and in P1’ at the C-terminal cleavage site. (B) PSK1 precursor expressed in *E. coli*, and purified by NiNTA chromatography, ultrafiltration and size exclusion chromatography. Proteins were separated by SDS-PAGE and stained by Coomassie Brilliant Blue (CBB). (C) proPSK1 is processed by SBT3.8 *in vitro*. The recombinant precursor was incubated with purified SBT3.8 or a mock-purified control fraction for the time indicated. Proteins were separated on Tris-Tricine gels and stained by CBB. (D) proPSK1 processing depends on SBT3.8. Recombinant proPSK1 was treated with exudates of wild-type or *sbt3.8* loss-of-function plants for the time indicated, separated on Tris-Tricine gels and stained by CBB. (E). SBT3.8 cleaves proPSK Asp-dependently at the C-terminal site irrespective of pH. Peptides released by SBT3.8 or the mock-purified control from recombinant proPSK1 at pH 7 or 5.5 were identified by mass spectrometry (top). No peptides were identified from the region of interest when Asp in P1’ was substituted by alanine (bottom). Cleavage assays have been repeated at least three times with similar results.

SBT3.8 was expressed with a C-terminal hexa-His tag in tobacco (*Nicotiana benthamiana*) leaves and purified from cell wall extracts by affinity chromatography as described (Stührwohldt et al., 2020). C-terminally hexa-His-tagged proPSK1 was expressed in *E. coli* and purified by affinity chromatography, ultrafiltration, and size exclusion chromatography (Figure 4B). Cleavage of recombinant proPSK1 was detected by SDS-PAGE after incubation with SBT3.8. Degradation was much reduced in a control digest, in which SBT3.8 was replaced by a mock-purified tobacco cell wall extract (Figure 4C). To test whether SBT3.8 is required for proPSK1 processing *in planta*, we incubated recombinant proPSK1 with exudates of wild-type and *sbt3.8* mutant seedlings grown in submerged culture. Rapid (within 5 min) cleavage was observed for wild-type but not for *sbt3.8* exudates (Figure 4D) indicating that cleavage is mediated by SBT3.8, and that there are no redundant activities, at least not at this stage of seedling development.

In order to identify the cleavage site(s) within the PSK1 precursor, recombinant proPSK1 was digested with SBT3.8 or the mock-purified negative control, the peptide fraction was recovered by ultrafiltration, peptides were purified on C18 stage tips and analyzed by mass spectrometry. We identified a series of peptides all indicative of cleavage at the C-terminal processing site (Figure 4E). N-termini were somewhat variable and likely result from unspecific cleavage by contaminating tobacco protease(s) that are abundant in cell wall extracts (Grosse-Holz et al., 2018). These peptides were detected specifically only in the SBT3.8 digest, irrespective of pH (7 or 5.5), and not for the control reaction (Figure 4E). To further test the significance of aspartate for C-terminal cleavage by SBT3.8, the cleavage assay was repeated under the same conditions with a site-directed proPSK1 mutant in which this Asp residue was replaced by Ala. Mass spectrometry failed to identify any peptides derived from C-terminal cleavage, neither at pH 7 nor at pH 5.5 (Figure 4E). The data indicate that SBT3.8 cleaves the PSK1 precursor specifically at the C-terminal processing site and that this cleavage depends on Asp in P1’. To test whether SBT3.8 is able to cleave other aspartate-containing pro-peptides, we purified recombinant proRGF1 from *E. coli* and incubated it with exudates of wild-type or *sbt3.8* mutant seedlings (Supplemental Figure 2A and B). proRGF1 was cleaved by wild-type but not by *sbt3.8* exudates, indicating that SBT3.8 is required for the observed processing of proRGF1. A site-directed D104A mutant of proRGF1 was resistant to cleavage by SBT3.8 (Supplemental Figure 2B), thus confirming cleavage at D104 and Asp-specificity of processing.

To verify that Asp-specific cleavage is required for PSK formation *in vivo*, we used generated transgenic Arabidopsis plants overexpressing wild-type proPSK1 from rice in translational fusion with GFP, and a site-directed mutant in which the two flanking Asp residues had been replaced by Ala (proPSK1ox-DD/AA). The functionality of rice proPSK1-GFP in Arabidopsis was confirmed in bioassays based on the growth-promoting activity of PSK (Kutschmar et al., 2009; Stührwohldt et al., 2011). Root length was increased by more than 50% and hypocotyls were significantly longer in three independent transgenic lines expressing proPSK1-GFP compared to Col-0 wild-type plants (Figure 5A, B). If Asp-specific cleavage were required for PSK biogenesis, we would expect the Ala-substituted proPSK1 mutant to be inactive in this assay. However, root and hypocotyl lengths of three independent proPSK1-DD/AA-GFP lines was increased to similar levels as in plants expressing the proPSK1-GFP wild-type construct (Figure 5A, B), even though processing is impaired by the D-to-A cleavage site mutation (Figure 5C). To reconcile the requirement for Asp at the C-terminal processing site for PSK1 formation (Figures 4E and 5C) with the apparent activity of the D-to-A substituted proPSK1 mutant *in vivo* (Figure 5A, B), we compared the expression of endogenous *PSK* genes in the transgenic and in wild-type plants. Interestingly, *PSK1, PSK2* and *PSK3* were found to be upregulated in plants carrying the rice proPSK1 overexpression constructs (Figure 5A), with particularly strong induction of PSK1 in the proPSK1ox-DD/AA overexpressors. The observed increase in root and hypocotyl length in plants expressing proPSK1-DD/AA may thus be explained by the upregulation of *PSK* genes other than rice *PSK1*, and does not necessarily mean that proPSK1-DD/AA is processed *in vivo* to release the mature PSK peptide. Alternatively, the SBT3.8-resistant proPSK1-DD/AA mutant may be cleaved at a different site to produce a longer version of the PSK peptide that nonetheless retains (some of the) activity of the mature PSK pentapeptide.

**Figure 5:**
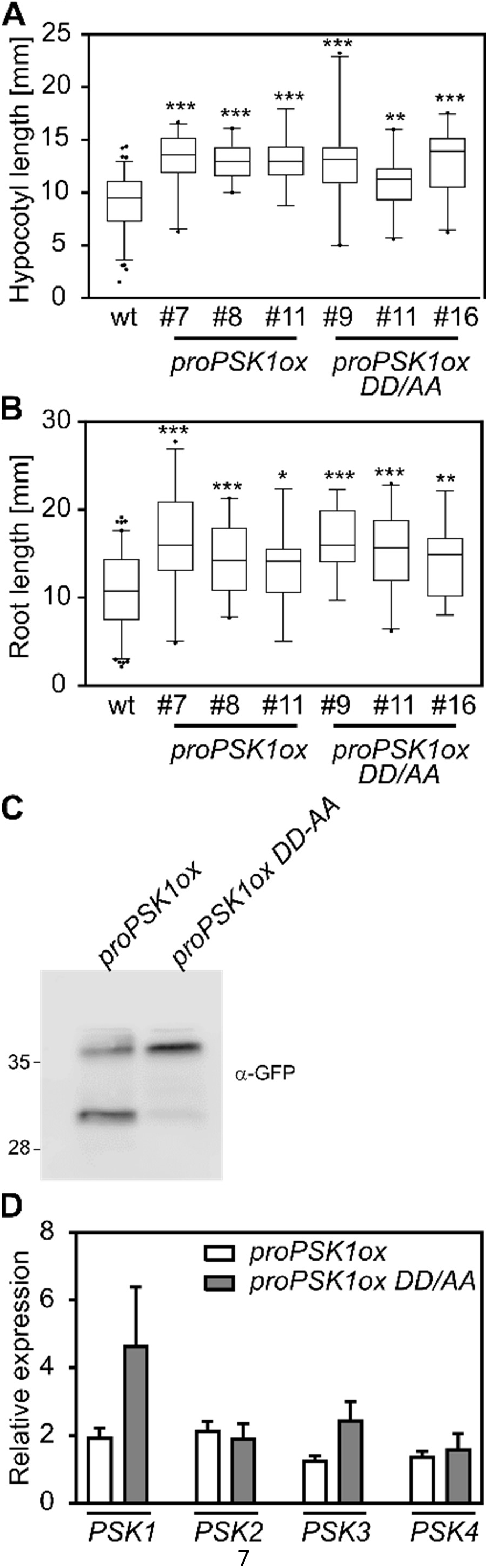
Root and hypocotyl growth are induced in plants expressing proPSK or the Asp-substituted proPSK mutant. (A-B) Hypocotyl length [mm] (A) and root length [mm] (B) of three independent lines overexpressing proPSK1-GFP or PSK1-DD/AA-GFP in comparison to the wild type (wt). Experiments were performed three times with similar results. Data are shown for one representative experiment as the mean ± SE ((A) n ≥ 16, (B) n ≥ 18). *, ** or *** indicate significant differences to the wild type at P < 0.05, P < 0.01 or P < 0.001 (two-tailed t test). (C) The DD/AA mutation impairs proPSK processing *in vivo*. Leaf protein extracts (20 μg) of representative PSK1ox-GFP or PSK1ox DD/AA-GFP lines were analyzed by anti-GFP immunoblotting. (D) qPCR analysis of plants overexpressing PSK1 or PSK1 DD/AA precursors. qPCR was performed on RNA isolated from 11 day-old plants in three biological replicates with two technical repeats. Gene regulation was normalized to three reference genes. Each time point included pooled plant material of several independent plants.

We further used the proPSK1ox lines to directly assess the effect of PSK on osmotic stress tolerance. Experimental plants were grown for five days on control media, then transferred to either control medium or mannitol-containing medium, and 6 days later, the increment of root growth was measured (Figure 6A). Root growth of wild-type plants was severely impaired on mannitol-containing media (Figure 6B), while proPSK1ox plants coped better with osmotic stress. The increment of root growth of proPSK1ox plants was about twice as large as in wild-type plants (Figure 6B). We further quantified the number of lateral roots and recorded significantly higher numbers of lateral roots in proPSK1ox lines. The data confirm a role for PSK in stress mitigation and, consistent with the upregulation of *PSK1, 3*, and *4* genes under stress (Figure 1), they indicate that the level of stress tolerance is limited by *PSK* expression levels.

**Figure 6:**
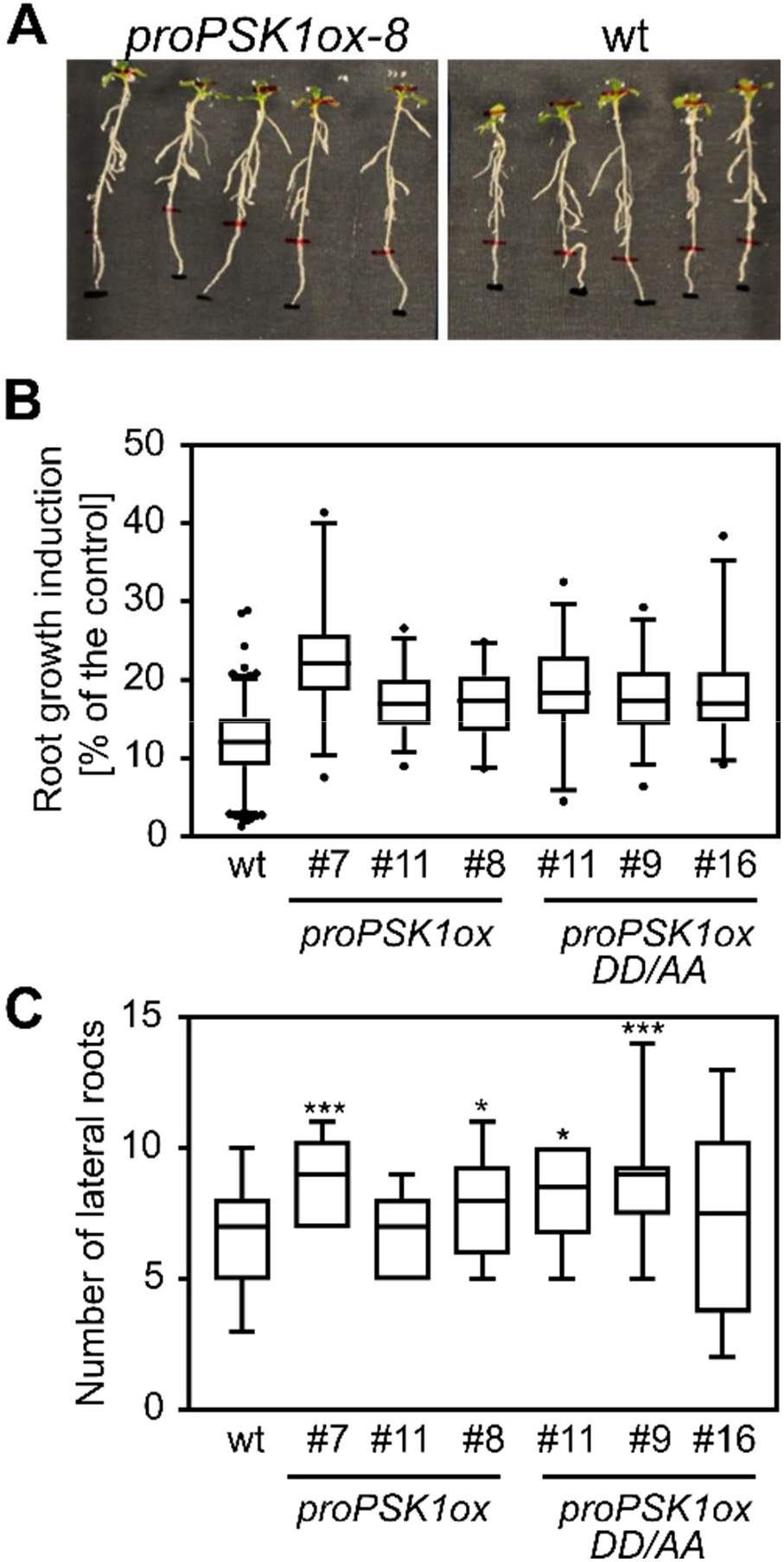
PSK1 overexpression improves adaptation to mannitol stress conditions. (A) Representative *proPSK1ox-8* and wild-type (wt) plants six days after transfer to mannitol-containing media. The increment of root growth is marked. (B) Root growth increment [mm] of wt and PSK overexpressing plants after transfer to mannitol-containing media shown as the percentage of plants grown on control media. (C) Number of lateral roots formed by overexpressors of proPSK1 or proPSK1-DD/AA compared to wt. Data are shown for one representative of three independent experiments as the mean ± SE ((B) n ≥ 10, (C) n ≥ 10). * and *** indicate significant differences to the wild type at P < 0.05 or P < 0.001, respectively (two-tailed t test) in comparison to the wild type.

The findings that *SBT* expression is induced upon osmotic stress (Figure 1), stress sensitivity is increased in the *sbt3.8* mutant (Figure 2B, C), and that SBT3.8 is able to process the PSK1 precursor (Figure 4), suggest that osmotic stress acclimation is regulated not only at the level of *PSK* expression, but also at the post-translational level of proPSK processing. To corroborate this notion and to confirm SBT3.8 as a limiting factor in stress mitigation, we generated transgenic Arabidopsis plants overexpressing SBT3.8 as a C-terminal sfGFP fusion. Overexpression of SBT3.8 was confirmed by qPCR and immunoblot analysis (Supplemental Figure S2A, B). Under control growth conditions, five-day-old SBT3.8ox seedlings had significantly longer main roots than wild type (Figure 7A), comparable to seedlings overexpressing proPSK1 (Figure 5B). On mannitol-containing media, SBT3.8ox plants showed increased osmotic stress tolerance compared to wild-type plants (Figure 7B-E). The number of lateral roots (Figure 7B), shoot and root fresh weight (Figure 7D and E) were all increased in SBT3.8ox compared to wild-type plants. SBT3.8ox plants also performed better than wild type when grown on soil under continuous drought stress (Supplemental Figure 2C). We finally analyzed the expression of genes involved in abscisic acid (ABA) biosynthesis including *ABA1* (zeaxanthin epoxidase), *ABA2* (cytosolic short-chain dehydrogenase/reductase forming ABA-aldehyde from xanthoxin) and AAO3 (aldehyde oxidase delta isoform catalyzing the final step in ABA biosynthesis). Under control conditions, the expression of *ABA1*, *ABA2* and *AAO3* was unchanged in SBT3.8 overexpressors compared to wild type (Figure 7F). Upon osmotic stress treatment, *ABA1, ABA2* and *AAO3* expression was induced to higher levels in *SBT3.8* overexpressors compared to wild type, although not all differences were statistically significant (Figure 7G). Increased ABA production may explain why *SBT3.8* overexpressors cope better with osmotic stress. The data support a regulatory role for SBT3.8 in the production of PSK as a signal for osmotic and drought stress acclimation.

**Figure 7:**
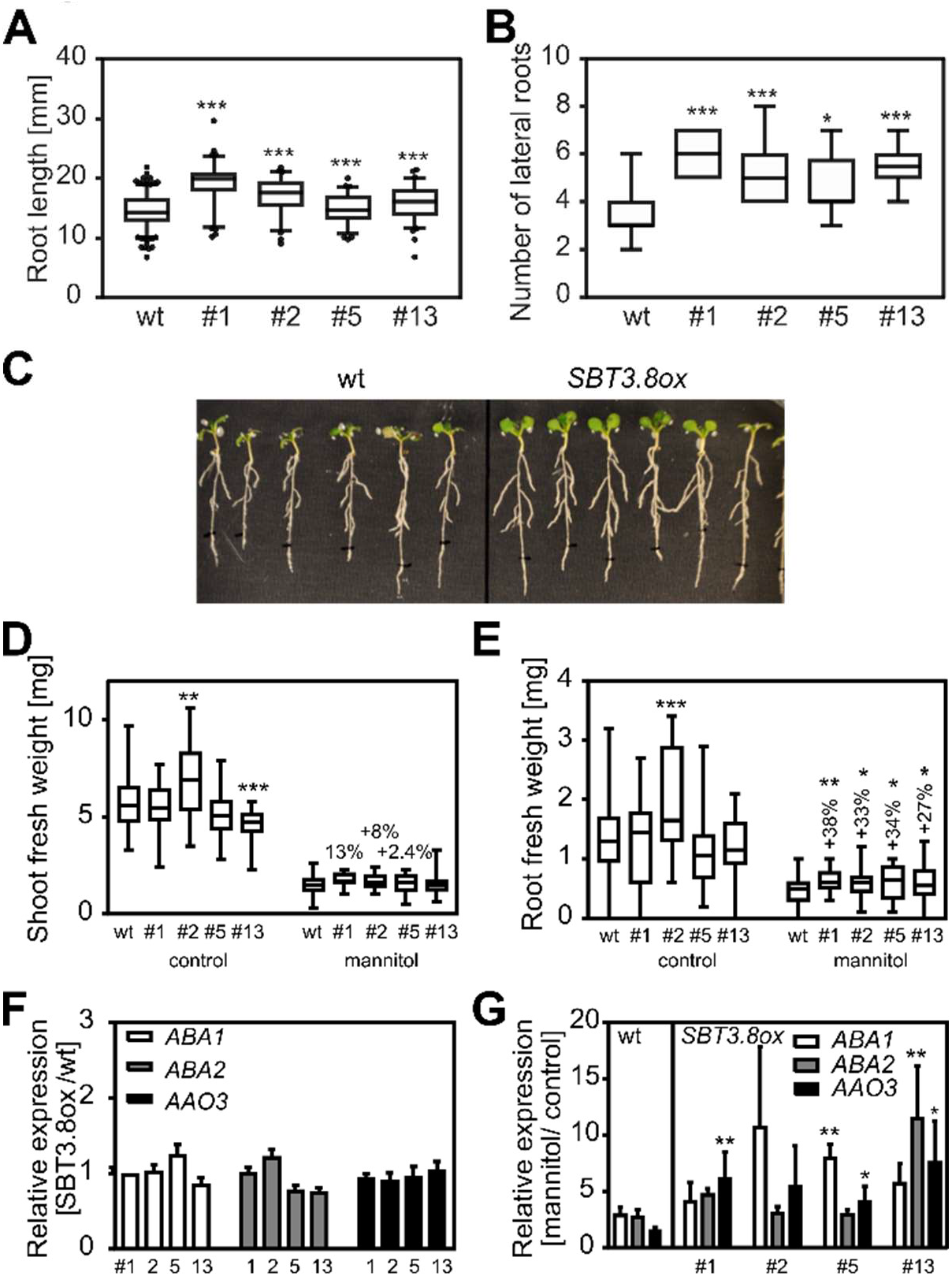
Constitutive SBT3.8 overexpression improves mannitol stress adaptation and induces growth. (A) Root length [mm] is significantly increased in four independent SBT3.8-overexpressing (*SBT3.8ox*) seedlings compared to wild type (wt). (B) Number of lateral roots, (C) shoot fresh weight [mg], and (D) root fresh weight [mg] of plants grown on mannitol-containing media are increased in *SBT3.8ox* compared to wt plants. Experiments were performed three times with similar results. Data are shown for one representative experiment as the mean ± SE ((A) n ≥ 66 (B) n ≥ 12, (C) n ≥ 30, (D) n ≥ 30). *, ** and *** indicate significant differences to the wild type at P < 0.05, P < 0.01 or P < 0.001, respectively (two-tailed t test). (F, G) Relative expression of *ABA1, ABA2* and *AAO3* in wt and *SBT3.8ox* lines under control (F) and osmotic stress conditions (G). Plants were grown for five days under control conditions and transferred to plates containing no or 300 mM of mannitol for six additional days. qPCR was performed on pooled plant material in three biological replicates with two technical repeats. Gene regulation was normalized to three reference genes. * or ** indicate significant differences to wt at P < 0.05 or P < 0.01, respectively (two-tailed t test).

## DISCUSSION

### PSK as a drought stress signal in Arabidopsis

PSK is known primarily for its growth regulatory properties. It was identified in 1996 as a mitogenic growth factor that stimulates cell division in Asparagus cell cultures (Matsubayashi and Sakagami, 1996). In Arabidopsis, PSK induces root and hypocotyl growth. The elongation of roots and hypocotyls is induced to similar levels by PSK treatment, by overexpression of the PSK1 precursor, and by overexpression of the PSK receptor PSKR1 (Kutschmar et al., 2009; Hartmann et al., 2013; Stührwohldt et al., 2011). These growth responses were found to rely on cell expansion rather than cell division (Stührwohldt et al., 2011; Kutschmar et al., 2009; Ladwig et al., 2015). However, in addition to its growth-regulatory properties, PSK is also involved in pollen germination and pollen tube elongation (Stührwohldt et al., 2015; Chen et al., 2000), vascular development (Holzwart et al., 2018), and in the regulation of plant immunity (Loivamäki et al., 2010; Igarashi et al., 2012; Mosher et al., 2013; Rodiuc et al., 2016; Zhang et al., 2018). Most recently, PSK was found to act as a peptide signal for stress-induced flower drop in tomato (Reichardt et al., 2020). Upon drought stress, PSK precursor genes and the precursor-processing subtilase SlPhyt2 are upregulated in the pedicel proximal to the abscission zone. Mature PSK acts in the abscission zone where it induced the expression of cell wall hydrolases to initiate the cell separation process (Reichardt et al., 2020). These findings prompted us to investigate whether PSK precursors and cognate subtilases may also be involved in drought stress responses in Arabidopsis.

We found that three subtilase genes (*SBT1.4, SBT3.7*, and *SBT3.8*) and three PSK precursor genes (*PSK1, PSK3*, and *PSK4*) are upregulated in Arabidopsis seedlings in response to osmotic stress. Tolerance to drought stress and osmotic stress was compromised in *sbt3.8* mutants and the latter could be restored by PSK treatment. In contrast, osmotic stress tolerance was improved in transgenic plants overexpressing either SBT3.8 or proPSK1. These data imply PSK signaling in drought stress mitigation in Arabidopsis, and they suggest SBT3.8 as a limiting factor in PSK formation.

### PSK precursor processing

The only protease previously implicated in PSK processing in Arabidopsis is SBT1.1. The protease was shown to cleave a synthetic peptide derived from the PSK4 precursor *in vitro* (Srivastava et al., 2008). SBT1.1-dependent processing of proPSK4 also was observed *in planta*, at least when the precursor was expressed ectopically under control of the constitutive CaMV 35S promoter. However, the SBT1.1 cleavage site is located several residues upstream of the PSK sequence and, therefore, cleavage by SBT1. 1 does not produce the mature and bioactive peptide (Srivastava et al., 2008). Also, no PSK-related phenotypes have been reported for the *sbt1.1* mutant (Srivastava et al., 2008). Therefore, the contribution of SBT1.1 to PSK maturation, if any, is questionable.

In tomato plants, PSK maturation depends on phytaspase-2 (SlPhyt2; (Reichardt et al., 2020)). Phytaspases were first identified in tobacco and rice (Chichkova et al., 2018). They constitute a subgroup of subtilases that cleave their substrates in an Asp-specific manner (hence the name: phyt<u>asp</u>ase is short for plant asp-specific protease). For substrate recognition and cleavage by phytaspases, Asp is required in the P1 position (immediately upstream of the scissile bond) (Chichkova et al., 2018; Reichardt et al., 2020). Consistently, SlPhyt2 cleaves PSK precursors at the amino-terminal processing site where the P1 Asp is conserved in all eight PSK precursors in tomato, and in the seven Arabidopsis PSK precursors as well (Reichardt et al., 2020; Kaufmann and Sauter, 2019).

In contrast to the twelve phytaspases tentatively identified in tomato (Reichardt et al., 2020), a single gene, *SBT3.8*, has been suggested as the phytaspase orthologue in Arabidopsis, cleaving substrate peptides specifically after Asp in P1 (Chichkova et al., 2018). Therefore, cleavage of PSK precursors by SBT3.8 would be expected at the N-terminus of the PSK peptide sequence. However, we observed processing of proPSK1 at the C-terminal site, which features Asp in the P1’ position (immediately downstream of the cleavage site), rather than in P1 (cf. Fig 4A for sequence of proPSK1). This Asp residue (D60 of proPSK1) turned out to be required for cleavage by SBT3.8 (Fig. 4D,E). Cleavage of proPSK1 at the N-terminal site was not observed, neither at physiological pH of 5.5, nor at pH 7.5 (Fig. 4E), at which SBT3.8 was reported to exhibit less stringent substrate specificity (Chichkova et al., 2018).

Cleavage of proPSK1 at D60 is fully consistent with our previous report on the biogenesis of CLEL6 and CLEL9. The respective precursors were found to be cleaved by SBT3.8 in an Asp-specific manner, and Asp was required in P1’ for processing (Stührwohldt et al., 2020). Likewise, we show here that cleavage of Root Growth Factor 1 (RGF1, alias CLEL8) by SBT3.8 depends on D104 (Supplementary Figure S1). In RGFs (alias CLEL peptides) the critical P1’ Asp residue marks the N-terminus of the mature peptides, in proPSK1 the Asp is located C-terminally to the PSK pentapeptide. We may thus anticipate a broader role for SBT3.8 in the biogenesis of peptide hormones carrying P1’ Asp at either the N– or the C-terminal processing site. With respect to PSK biogenesis, SBT3.8 appears to be specific for the PSK1 precursor, as this is the only one featuring Asp in P1’. Hence, other unknown proteases must be involved in C-terminal processing of Arabidopsis proPSK2 to 7. SBTs 1.4 and 3.7 are obvious candidates, as their expression is upregulated in response to osmotic stress as well.

The protease(s) responsible for N-terminal processing of Arabidopsis PSK precursors also remain(s) to be identified. Similar to PSK maturation in tomato (Reichardt et al., 2020), a *bona fide* Arabidopsis phytaspase may be involved in this processing step. Alternatively, the introduction of a negative charge in P1’ by tyrosine sulfation may allow recognition of the N-terminal site by SBT3.8. Tyrosine sulfation is known to be required for full peptide activity (Matsubayashi et al., 2006; Kutschmar et al., 2009). Mutants impaired in tyrosylprotein sulfotransferase activity show severe developmental defects that can be rescued by addition of the mature di-sulfated PSK peptide (Kutschmar et al., 2009; Stührwohldt et al., 2011; Matsubayashi et al., 2006). The requirement of tyrosine sulfation for bioactivity was rationalized by structural studies showing that the two sulfate moieties contribute to the interaction of PSK with its receptor PSKR1, thus sensitizing PSK perception by PSKR1 (Wang et al., 2015). The hypothetical contribution of sulfo-tyrosine to cleavage site recognition and peptide biogenesis provides another explanation for the strong phenotype observed in tyrosylprotein sulfotransferase loss-of-function mutants.

### Complex regulation of osmotic stress responses by peptides

In addition to PSK, other peptides are known to contribute to plant responses to osmotic stress. A cysteine-rich peptide in rice, OsDT11, enhances drought tolerance in rice in an abscisic acid (ABA)-dependent manner (Li et al., 2017). Transgenic plants overexpressing OsDT11 show enhanced drought tolerance, reduced water loss, reduced stomatal density and increased levels of ABA (Li et al., 2017). PROPEPs are primarily known as precursors of immune-regulatory phytocytokines (Huffaker and Ryan, 2007). However, one family member, Arabidopsis *PROPEP3*, was recently shown to be involved in salt stress responses (Nakaminami et al., 2018). *AtPROPEP3* expression increases in response to salt stress, and salt tolerance correlates with *AtPROPEP3* expression levels in transgenic plants. Stress tolerance is mediated by a fragment of the PEP3 phytocytokine and depends on the PEP receptor PEPR1 (Nakaminami et al., 2018). Likewise, CAP-derived peptide (CAPE) also was first identified as an immune-modulatory peptide which is derived from the C-terminus of tomato pathogenesis-related protein 1 (PR1; (Chen et al., 2014). In Arabidopsis, CAPE1 negatively regulates salt tolerance (Chien et al., 2015). Also in Arabidopsis, drought stress was shown to induce the production of CLE25 as a long-distance signal for increased resistance to dehydration. CLE25 binds to the BAM3 receptor in the leaves and regulates stomatal aperture in an ABA-dependent manner (Takahashi et al., 2018). Here we add PSK to the suite of drought-stress signals in Arabidopsis, and SBT3.8 as a limiting factor for PSK production. Which proteases and the extent to which they contribute to the production of the other peptides remains to be seen. Their number may well exceed the number of peptides they produce, thus highlighting the complexity of peptide-mediated osmotic stress responses.

## METHODS

### Plant Materials and Growth Conditions

For growth experiments on synthetic media, Arabidopsis seeds were surface-sterilized in 70% (v/v) ethanol for 15 minutes, washed in 100% ethanol and laid out on square plates containing 0.5 x MS (Murashige and Skoog, 1962), 1% (w/v) sucrose and 0.38% (w/v) gelrite. Seeds were stratified for one day at 4° C and grown under short-day conditions (12 h photoperiod) at 22 °C and 100-120 μE white light. For osmotic stress treatment, 5-day old seedlings were transferred to plates containing the same medium supplemented with 300 mM mannitol. Growth phenotypes were scored after 6 days and compared for plants grown on control and mannitol-containing media, respectively. Root length was measured using ImageJ (http://rsbweb.nih.gov/ij/). All experiments were carried out at least three times; representative results are shown. PSK peptide was added at indicated concentrations. The peptide was synthesized by PepMIC (Souzhou, China) and obtained at > 95% purity. Lyophilized peptides were reconstituted in water. Peptide concentration was determined by a modified Lowry procedure using the DC Protein Assay (BioRad; Munich, Germany) according to the manufacturer’s instructions. Seeds of *N. benthamiana* were provided from Agroscience (Agroscience GmbH, Neustadt, Germany) and were grown at a day-length of 16 hours at 28 °C. Tobacco plants were fertilized 3 times a week with universal fertilizer (Wuxal®, 2ml/l) and used for experiments at the age of four to six weeks. The *sbt* loss-of-function mutants and the SBT3.8ox transgenic lines have been described before (Rautengarten et al., 2005; Stührwohldt et al., 2020). For experiments on soil, Arabidopsis plants were grown on potting compost with 3.6 % (v/v) sand and 7.2 % (v/v) perlite. To minimize the risk of thrips infection, seeds were incubated over night at −75 °C. They were re-suspended in 0.1% (w/v) agar, stratified for 48 hours at 4 °C in the dark, and grown under short-day conditions (12 h photoperiod) at 22 °C and 100-120 μE white light. Plants were watered as indicated. For drought stress experiments, wt and *SBT3.8ox* plants were grown for three weeks with sufficient water. Subsequently, plants were watered only every third day with 10 ml each.

### Generation of Expression Constructs

All primers used for amplification of constructs by PCR are listed in Supplemental Table 1. ORFs of PSK1 and RGF1 lacking the predicted signal peptides were amplified by PCR from cDNA and cloned into the *NcoI* restriction site of pETDuet1 (Novagen/Merck KGaA, Darmstadt, Germany). C-terminal His tags were added by including 6 His codons in the reverse PCR primers. Correct orientation and translational fusion with the C-terminal His-tag were verified by sequencing. PSK1 and RGF1 D/A mutants were created by site-directed mutagenesis or PCR with reverse primer including mutations, respectively. Mutations were confirmed by sequencing.

### Expression and Purification of Recombinant Proteins

*Escherichia coli* cells were grown in LB medium with appropriate antibiotics at 220 rpm and 37 °C to an A600 of 0.6. Protein expression in *E. coli* BL21 was induced by 1 mM IPTG for two hours at 30 °C. His-tagged PSK1 or RGF1 precursors were purified from bacterial extracts by metal chelate affinity chromatography on NiNTA Agarose (Qiagen, Hilden, Germany) according to the manufacturer’s recommendations. Cells were harvested by centrifugation (4000 x g, 4 °C, 20 min) and resuspended in 50 mM sodium phosphate buffer, pH 7.0, supplemented with 300 mM NaCl including lysozyme, PMSF, DNAse and Proteinase Inhibitor Cocktail (Serva). Cells were lysed by sonication (SONOPULS HB2070 with MS72 sonic needle, Bandelin Electronics; Berlin, Germany) thrice at 4 °C for 30 seconds. The soluble fraction was collected by centrifugation (15000 x g, 4 °C, 10 min). Recombinant proteins were bound to the Ni-NTA column for at least one hour. The column was washed extensively in buffer containing 10 mM imidazole and recombinant proteins were eluted in buffer containing 200 mM imidazole. After ultrafiltration (30 kDa molecular mass cutoff), the recombinant protein in the filtrate was further purified by size exclusion chromatography (NGC™ chromatography system with Enrich™ SEC 650 column, BioRad, Munich, Germany) in 50 mM NaH2PO4 /Na2HPO4, pH 5.5 (or 7.0 if indicated), 10 mM NaCl.

### Generation of PSK1 Overexpressors

The cDNAs of rice (*Oryza sativa) PSK1* and the site-directed D79A, D85A double mutant were a gift of Andrey Vartapetian (Moscow State University). They were obtained in translational fusion with EGFP (enhanced green fluorescent protein) between the CaMV 35S promoter and terminator in pRTL2 (Carrington et al., 1990).The entire expression cassettes were cut out from pRTL2 with PstI, and cloned into pGreenII 0229. Plasmids were transformed into GV3101 and transformed into *Arabidopsis thaliana* Col-0 by floral dip (Clough and Bent, 1998). Transgenic lines were selected by glufosinate and homozygous lines in the T3 or T4 generation were used in further experiments.

### Transient Expression in *N. benthamiana* and Protein Extraction

*Agrobacterium tumefaciens* strains C58C1 and GV3101 were used for transient expression experiments in *N. benthamiana*. Bacteria were grown on plates containing appropriate antibiotics (rifampicin, tetracycline and spectinomycin for C58C1, and gentamycin and spectinomycin for GV3101, respectively) at 28 °C and were washed off the plates in 10 mM MES, pH 5.6 containing 10 mM MgCl2. Prior to infiltration, 150 μM acetosyringone were added to the bacterial suspension. A blunt syringe was used to infiltrate the suspension into the leaves. Five days after infiltration, the leaves were harvested. For collection of apoplastic fractions, the harvested leaf material was washed three times with cold ddH2O and stored in ice-cold infiltration buffer (300 mM NaCl, 50 mM sodium phosphate buffer pH 7.0) and vacuum-infiltrated twice at 70 mbar for one minute. Subsequently, tobacco leaves were blotted dry and transferred to a syringe barrel filled with glass wool and centrifuged for ten minutes at 4°C and 950 x g. The resulting protein solution was centrifuged at 16.000 x g for additional ten minutes. His-tagged SBT3.8 was purified from the supernatant on Ni-NTA agarose as described above. SBT3.8 was bound to the column for two hours, the column was washed extensively in buffer containing 20 mM imidazole and SBT3.8 was eluted in buffer containing 400 mM imidazole.

### Cleavage of Peptide Precursors by SBT3.8 and Identification of Cleavage Products

Approximately 10 μg of recombinant pro-peptide and 100 ng of SBT3.8, or the same volume of the mock-purified fraction from empty-vector infiltrated plants as control, were incubated at room temperature for 30 minutes in a total volume of 30 μl of 50 mM sodium phosphate buffer, pH 7.0 or 5.5, supplemented with 10 mM NaCl. For subsequent MS analysis, the reaction was stopped at 95 °C for 10 minutes. If indicated, peptides were separated by ultrafiltration with a molecular mass cutoff of 10 kDa. Peptides were purified on C18 reverse phase stage tips (Agilent, Santa Clara, California, USA). C18 discs were conditioned in 80% (v/v) acetonitrile (ACN) and equilibrated in 0.5% (v/v) acetic acid by centrifugation at 5000 rpm for 1 min each. Peptides were bound by centrifugation at 3000 rpm for 1 min, washed in 0.5% (v/v) acetic acid for 1 min and eluted by 50% (v/v) ACN. The isolated peptides were dried by vacuum evaporation. For subsequent SDS-PAGE analysis, the reaction was stopped by incubation at 95 °C for 10 minutes including loading dye. Samples were separated by SDS-PAGE or Tris-Tricine PAGE (Schägger, 2006). Wild-type and *sbt3.8* mutant exudates were obtained by growing seedlings in submerged culture in 0.5 x MS medium with 1% (w/v) sucrose for ten days as described (Ohyama et al., 2008). Under these conditions, seedlings release their extracellular protein content into the medium (Ohyama et al., 2008). Apoplastic proteins were enriched by ultrafiltration with a molecular mass cutoff of 5 kDa. Exudates were used at a protein equivalent of 500 ng for cleavage assays.

### Immunodetection

For western blots, proteins were transferred to nitrocellulose membranes using a semi-dry blotter and standard protocols (BioRad). GFP-fusion proteins were detected by polyclonal α-GFP antiserum (Thermo Fisher Scientific, Waltham, Massachusetts, USA), SBTs by a mixture of polyclonal antisera against tomato SBTs 1 to 4, His_6_-tagged proteins by a monoclonal mouse α–His_6_ antibody from Dianova (Hamburg, Germany), and Flag-tagged proteins by α–Flag antibody directly coupled to horseradish peroxidase (Taufkirchen, Germany) at the indicated concentrations. Except for α–Flag, horseradish peroxidase-conjugated anti-rabbit or anti-mouse IgG was used as secondary antibody, followed by enhanced chemiluminescence detection using a Li-Cor Odyssee imager (Li-Cor Biosciences, Bad Homburg, Germany).

### Mass Spectrometry

Peptides were analyzed via LC-MS/MS using the nanoflow EASY-nLC 1000 System (Thermo Fisher Scientific) for HPLC separation and an Orbitrap hybrid mass spectrometer (Q-Exactive, Thermo Scientific; Waltham, Massachusetts, USA) for mass analysis. Peptides were eluted from a 75 μm analytical C18 column (PepMan, Thermo Fisher Scientific) on a linear gradient from 4% (v/v) to 64% (v/v) ACN over 135 min, and sprayed directly into the Q-Exactive Plus mass spectrometer. Proteins were identified via MS/MS on basis of the fragmentation spectra of multiple charged peptides. Up to twelve data-dependent MS/MS spectra were acquired in the Orbitrap for each full-scan spectrum. Full-scan spectra were acquired at 70,000 full-width half-maximum resolution, fragment spectra at a resolution of 35,000 full-width half-maximum, respectively. An inclusion list was used for fragmentation containing peptides covering the C-terminus of the precursor which includes the mature PSK pentapeptide.

### Real-time PCR

RNA isolation was performed as previously described with minor modifications (Kutschmar et al., 2009). cDNA was synthesized using up to 5 μg of total RNA, oligo dT primers and RevertAid Reverse Transcriptase (Thermo Fisher Scientific). SBT primers for quantitative real-time PCR analysis were described and used previously (Stührwohldt et al., 2020). Quantitative PCRs (total volume 25 μl) were performed in biological triplicates with two technical repeats on the obtained cDNAs using a CFX96 Real-Time PCR Detection system (BioRad). Primer efficiencies and optimal primer concentrations were determined experimentally. qPCR was performed with Taq polymerase expressed in and purified from *E. coli* and SYBR-Green (Cambrex Bio Science Rockland Inc.; Rockland, ME, USA). Relative mRNA levels were determined after normalization to three reference genes (*Actin2, EF* and *Tubulin*) following the equation by (Pfaffl, 2001).

## Author Contributions

NS and EB performed experiments; NS and AS conceived the project; MS provided preliminary data and conceptual input; NS and AS designed the research strategy; NS and AS supervised the research and wrote the paper.

## Acknowledgements

Authors thank Lhana Stein, Isolde Schuck (both Department of Plant Physiology and Biochemistry, Institute of Biology, University of Hohenheim) and Timo Staffel (Plant Developmental Biology and Physiology, University of Kiel) for excellent technical support. We also thank Andrey Vartapetian (Moscow State University) for the proPSKox-GFP and proPSKox-DD/AA-GFP overexpression constructs, and Jens Pfannstiel and Berit Würtz at the Hohenheim University’s Core Facility Service Unit Mass Spectrometry for mass spectral analyses.

---

**Supplemental Figure S1**: **RGF1 is cleaved by SBT3.8 in an Asp-dependent manner** (A) Amino acid sequence of proRGF1. The mature RGF1 peptide is underlined. The aspartate residue at the N-terminal cleavage site is highlighted in red. (B,C) The RGF1 wild-type precursor (B) and its site-directed D104A mutant (C) were expressed and purified from *E. coli*. Precursor proteins were incubated with exudates from wild-type or *sbt3.8* mutant seedlings for the time indicated. Digests were separated on Tris-Tricine gels and stained by CBB. The cleavage product generated from the RGF1 precursor by wild-type exudates is indicated (arrow).

**Supplemental Figure 2: Characterization of *SBT3.8ox* lines with respect to SBT3.8 transcript and protein levels, and drought stress tolerance** (A) Relative expression of *SBT3.8* in four independent P35S:SBT3.8-sfGFP (*SBT3.8ox*) lines under normal growth conditions in comparison to the wild type (wt). (B) SBT3.8-sfGFP protein levels in four independent *SBT3.8ox* lines in comparison to the wt. Leaf protein extracts (20 μg) were separated by SDS-PAGE and analyzed on anti-GFP immunoblots. (C) Representative wt and *SBT3.8ox* plants are shown that were grown for three weeks with sufficient water. Subsequently, plants were watered only every third day with 10 ml each. Pictures were taken after three weeks of water deficit.

